# Depletion of Abundant Sequences by Hybridization (DASH): Using Cas9 to remove unwanted high-abundance species in sequencing libraries and molecular counting applications

**DOI:** 10.1101/031708

**Authors:** W Gu, ED Crawford, BD O’Donovan, MR Wilson, ED Chow, H Retallack, JL DeRisi

**Affiliations:** Departments of Pathology and Laboratory Medicine, University of California, San Francisco; Department of Biochemistry and Biophysics, University of California, San Francisco; Howard Hughes Medical Institute; Integrative Program in Quantitative Biology, Bioinformatics, University of California San Francisco; Department of Neurology, University of California, San Francisco; Center for Advanced Technology, Department of Biochemistry and Biophysics, University of California, San Francisco; Medical Scientist Training Program, Biomedical Sciences Graduate Program, University of California, San Francisco

**Keywords:** Cas9, CRISPR, Depletion, Sequencing, RNA-Seq, KRAS, Cancer, diagnostics

## Abstract

* Wei Gu and Emily Crawford contributed equally to this work

**Background:** With widespread adoption of next-generation sequencing (NGS) technologies, the need has arisen for a broadly applicable method to remove unwanted high-abundance species prior to sequencing. We introduce DASH (Depletion of Abundant Sequences by Hybridization), a facile technique for targeted depletion of undesired sequences.

**Results**: Sequencing libraries are DASHed with recombinant Cas9 protein complexed with a library of single guide RNAs (sgRNAs) programmed to target unwanted species for cleavage, thus preventing them from consuming sequencing space. We demonstrate up to 99% reduction of mitochondrial ribosomal RNA (rRNA) in HeLa cells, and enrichment of pathogen sequences up to 4-fold in metagenomic samples from patients with infectious diseases. Similarly, we demonstrate the utility of DASH in the context of cancer diagnostics by significantly increasing the detectable fraction of *KRAS* mutant sequences over the predominant wild-type allele.

**Conclusion:** This simple single-tube method is reprogrammable for virtually any sample type to increase sequencing yield without additional cost.

## Background

The challenge of extracting faint signals from abundant noise in molecular diagnostics is a recurring theme across a broad range of applications. In the case of RNA sequencing (RNA-Seq) experiments specifically, there may be several orders of magnitude difference between the most abundant species and the least. This is especially true for metagenomic analyses of clinical samples like cerebrospinal fluid (CSF), whose source material is inherently limited [1], making enrichment or depletion strategies impractical or impossible to employ prior to library construction. The presence of unwanted high-abundance species, such as transcripts for the 12S and 16S mitochondrial ribosomal RNAs (rRNAs), effectively increases the cost and decreases the sensitivity of counting-based methodologies. The same issue affects other molecular clinical diagnostics. In cancer profiling, the fraction of the mutant tumor-derived species may be vastly outnumbered by wild-type species due to the abundance of immune cells or the interspersed nature of some tumors throughout normal tissue. This problem is profoundly exaggerated in the case of cell-free DNA/RNA diagnostics, whether from malignant [2, 3], transplant [4], or fetal sources [5, 6], and relies on brute force counting by either sequencing or digital PCR (dPCR) [7] to yield a detectable signal. For these applications, a technique to deplete specific unwanted sequences that is independent of sample preparation protocols and agnostic to measurement technology is highly desired.

CRISPR (Clustered Regularly Interspaced Short Palindromic Repeats) and Cas (CRISPR associated)-nucleases, such as Cas9, function in bacterial adaptive immune systems to remove incoming phage DNA from the host without harm to the bacteria’s own genome. The CRISPR-Cas9 system has attained widespread adoption as a genome editing technique [8–11]. When coupled with single guide RNAs (sgRNAs) designed against targets of interest, S. *pyogenes* Cas9 binds to 3’ NGG protospacer adjacent motif (PAM) sites and produces double stranded breaks if the sgRNA successfully hybridizes with the adjacent target sequence (Figure 1A). *In vitro,* Cas9 may be used to cut DNA directly, in a manner analogous to a conventional restriction enzyme, except that the target sequence (outside of the PAM site) may be programmed at will and massively multiplexed without significant off-target effects. This affords the unique opportunity to target and prevent amplification of undesired sequences, such as those that are generated during next-generation sequencing (NGS) protocols.

**Figure 1.**
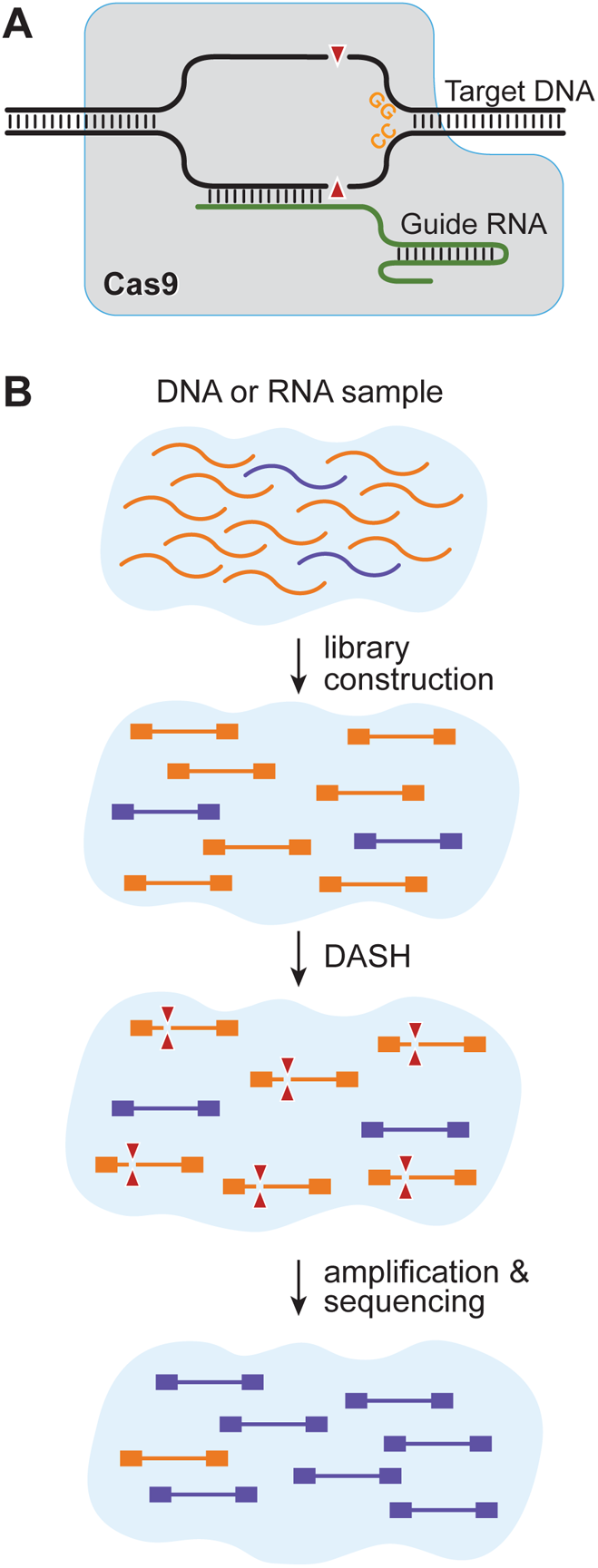
A) S. *pyogenes* Cas9 protein binds specifically to DNA targets that match the ‘NGG’ protospacer adjacent motif (PAM) site. Additional sequence specificity is conferred by a single guide RNA (sgRNA) with a 20 nucleotide hybridization domain. DNA double strand cleavage occurs three nucleotides upstream of the PAM site. B) Depletion of Abundant Sequences by Hybridization (DASH) is used to target regions that are present at a disproportionately high copy number in a given next-generation sequencing library following tagmentation or flanking sequencing adaptor placement. Only non-targeted regions that have intact adaptors on both ends of the same molecule are subsequently amplified and represented in the final sequencing library.

In this paper, we have exploited the unique properties of Cas9 to selectively deplete unwanted high-abundance sequences from existing RNA-Seq libraries. We refer to this approach as Depletion of Abundant Sequences by Hybridization (DASH). Employing DASH after transposon-mediated fragmentation but prior to the following amplification step (which relies on the presence of adaptor sequences on both ends of the fragment) prevents amplification of the targeted sequences, thus ensuring they are not represented in the final sequencing library (Figure 1B). We show that this technique preserves the representational integrity of the non-targeted sequences while increasing overall sensitivity in cell line samples and human metagenomic patient samples. Further, we demonstrate the utility of this system in the context of cancer detection, in which depletion of wild-type sequences increases the detection limit for oncogenic mutant sequences. The DASH technique may be used to deplete specific unwanted sequences from existing Illumina sequencing libraries, PCR amplicon libraries, plasmid collections, phage libraries, and virtually any other existing collection of DNA species.

Existing specific sequence enrichment techniques - such as pull-down methods [5, 12-14], amplicon-based methods [3, 15], molecular inversion methods (Turner et al 2009, Akhras et al 2007, Hiatt et al 2013), COLD-PCR [16], Competitive Allele-Specific TaqMan PCR (castPCR) [17], and the classic method of using restriction enzyme digestion on mutant sites [18] - can effectively enrich for targets in sequencing libraries, but these are not useful for discovery of unknown or unpredicted sequences. Brute force counting methods also exist, such as digital PCR (Pan et al., 2015; Vogelstein and Kinzler, 1999), but they are not easy to multiplex across a large panel. While high-throughput sequencing of select regions can be highly multiplexed to detect rare and novel mutations, and barcoded unique identifiers can overcome sequencing error noise [19], it is costly since the vast majority of the sequencing reads map to non-informative wild-type sequences. A number of sequence-specific RNA depletion methods also currently exist. Illumina’s Ribo-Zero rRNA Removal Kit and Ambion’s GLOBINclear Kit pull rRNAs and globin mRNAs, respectively, out of total RNA samples using sequence-specific oligos conjugated to magnetic beads. RNAse H-based methods, such as New England BioLab’s NEBNext rRNA Depletion Kit similarly mark abundant RNA species with sequence-specific DNA oligos, and then subject them to degradation by RNAse H, which digests RNA/DNA hybrid molecules [20]. These methods are all employed prior to the start of library prep, and are limited to samples containing at least 10-100 ng of RNA. DASH, in contrast, depletes abundant species after complementary DNA (cDNA) amplification, and thus can be utilized for essentially any amount of input sample.

## Results

We demonstrate deletion of unwanted mitochondrial ribosomal RNA using DASH first on HeLa cell line RNA (Figure 2A,B) and then on CSF RNA from patients with pathogens in their CSF, in order to increase sequencing bandwidth of useful data. Selection of rRNA sgRNA targets was based on examining coverage plots for standard RNA-Seq experiments on HeLa cells as well as on several patient CSF samples. Fifty-four sgRNA target sites within this region of the mitochondrial chromosome were chosen, situated approximately every 50 bp over a 2.5 kb region (sequences listed in Supplemental Table 1). sgRNA sites are indicated by red arrows in Figure 2B. sgRNAs for these sites were generated as described in the methods section. Coverage of the 12S and 16S mitochondrial rRNA genes was consistently several orders of magnitude higher than the rest of the mitochondrial and non-mitochondrial genes (Figures 2C and 3).

**Table 1:** Summary of depletion/enrichment results in DASH-treated clinical CSF samples.

|  |  |  | targeted genes (fpkm) |  |  |  | representative pathogenic gene* (alignment rate) |  | R<sup>2</sup> |
| --- | --- | --- | --- | --- | --- | --- | --- | --- | --- |
| Pathogen | library size (reads) |  | 12S |  | 16S |  |  |  | non-targeted genes untreated vs DASHed |
|  | untreated | DASHed | untreated | DASHed | untreated | DASHed | untreated | DASHed |  |
| *B. mandrillaris* | 1,813,373 | 2,537,356 | 298,922 | 28,005 | 380,073 | 93,164 | 0.028% | 0.102% | 0.992 |
| *C. neoformans* | 2,947,193 | 3,425,784 | 361,501 | 37,164 | 342,857 | 93,703 | 1.5% | 15.4% | 0.986 |
| *T. solium* | 2,382,766 | 1,890,534 | 451,044 | 46,993 | 317,640 | 43,257 | 12.0% | 44.3% | 0.994 |

\* Representative genes are 16S for *B. mandriallaris* and 18S for *C. neoformans* and *T. solium*

**Figure 2.**
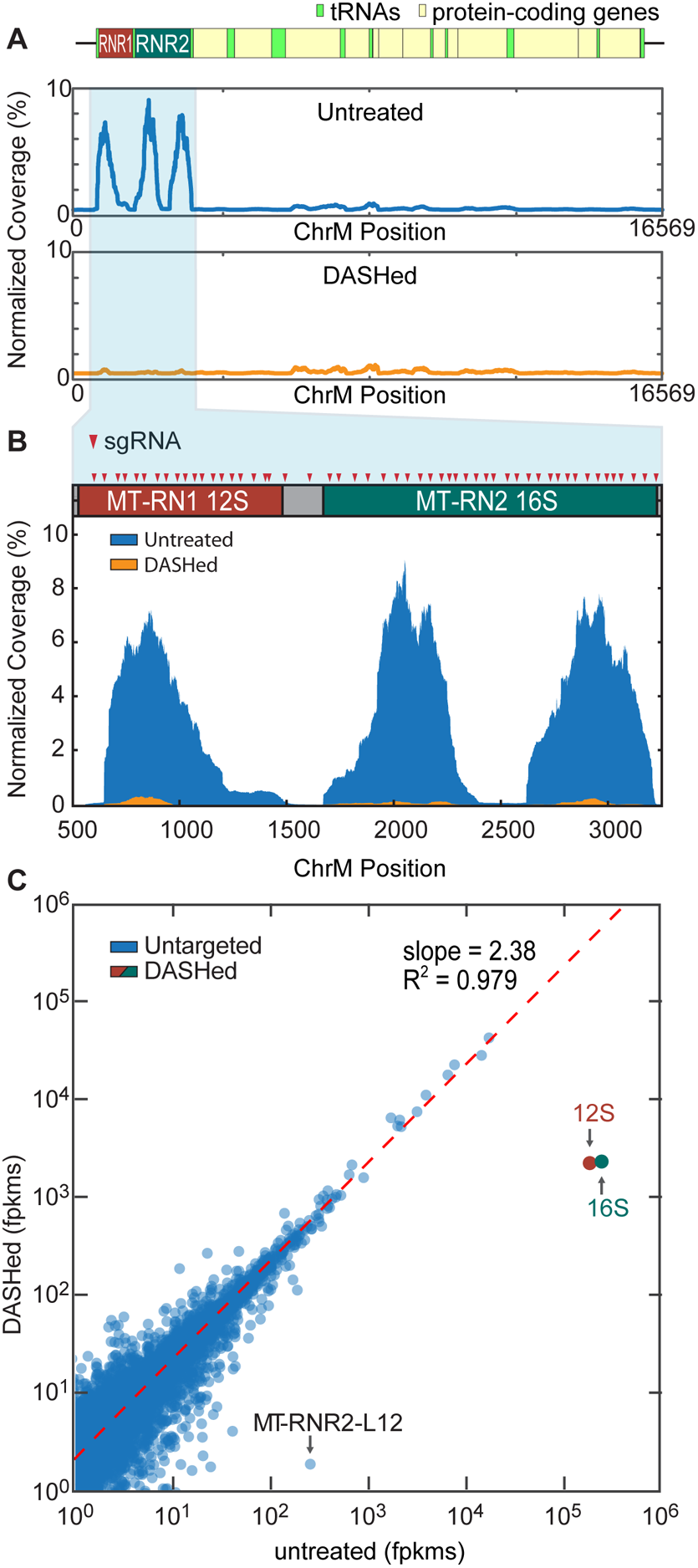
Depletion of Abundant Sequences by Hybridization (DASH) targeting abundant mitochondrial ribosomal RNA in HeLa RNA extractions. A) Normalized coverage plots showing alignment to the full-length human mitochondrial chromosome. Before treatment, three distinct peaks representing the ribosomal 12S and 16S subunits characteristically account for a large majority of the coverage (>60% of total mapped reads). After treatment, the peaks are virtually eliminated – with 12S and 16S signatures reduced 100-fold to 0.055% of mapped reads. B.) Coverage plot of Cas9-targeted region with 12S and 16S gene boundaries across the top. Each red triangle represents one sgRNA target site. 54 target sites were chosen, spaced approximately 50 bp apart. C) Scatterplot of the log of fragments per kilobase of transcript per million mapped reads (log-fpkm) values per human gene in the control vs. treated samples illustrate the significant reduction in reads mapping to the targeted 12S and 16S genes. DASH treatment results in 82 and 105-fold reductions in coverage for the 12S and 16S subunits, respectively. The slope of the regression line (red) fit to the untargeted genes indicates a 2.9-fold enrichment in reads mapped to untargeted transcripts. R-squared (R^2^) value of the regression line (0.979) indicates minimal off-target depletion. Between replicates, the R^2^ coefficient between fpkm values across all genes is 0.994, indicating high reproducibility (three replicates). Notably, one gene,MT-RNR2-L12 (MT-RNR2-like pseudogene), shows significant depletion in the DASHed samples compared to the control.

To calculate the input ratio of Cas9 and sgRNA to sample nucleic acid, we estimated that 90% of each sample was comprised of the rRNA regions that we targeted, thus our potential substrate makes up 4.5 ng of a 5 ng sample. This corresponds to a target site concentration of 13.8 nM in the 10 μL reaction volume. To assure the most thorough Cas9 activity possible, and given that Cas9 is a single-turnover enzyme *in vitro* [21], we used a 100-fold excess of Cas9 protein and a 1,000-fold excess of sgRNA relative to the target. Thus, each 10 μL sample of cDNA generated from a CSF sample contained a final concentration of 1.38 μM Cas9 protein and 13.8 μM sgRNA. In the case of HeLa cDNA, we used only 1 ng per sample, and therefore decreased the Cas9 and sgRNA concentrations by 5-fold. However, since mitochondrial rRNA sequences represented only 60% of the HeLa samples (compared to 90% for CSF), the HeLa samples contained 150-fold Cas9 and 1,500-fold sgRNA. To examine dose response, we processed additional 1 ng HeLa samples treated with 15-fold Cas9 and 150-fold sgRNA. Both concentrations were done in triplicate (see Supplemental Figure 1).

### Reduction of Unwanted Abundant Sequences in HeLa Samples

We first demonstrate the utility and efficacy of our approach using sequencing libraries prepared from total RNA extracted from HeLa cells. In the untreated samples, reads mapping to 12S and 16S mitochondrial rRNA genes represent 61% of all uniquely-mapped human reads. After DASH treatment, these sequences are reduced to only 0.055% of those reads (Figure 2A and B). Comparison of gene-specific fragments per kilobase of transcript per million mapped reads (fpkm) values between treated and untreated samples reveals mean 82-fold and 105-fold decreases in fpkms for 12S and 16S rRNA, respectively, in the samples treated with 150-fold Cas9 and 1,500-fold sgRNA (Figure 2C). Similarly, the samples treated with 10-fold Cas9 and 100-fold sgRNA show 30 and 45-fold reductions in 12S and 16S fpkm values, respectively, indicating a dose-dependent response to DASH treatment (Supplemental Figure 1).

### Enrichment of Non-Targeted Sequences and Analysis of Off-Target Effects in HeLa Samples

This profound depletion of abundant 12S and 16S transcripts increases the available sequencing capacity for the remaining, untargeted transcripts. We quantify this increase by the slope of the regression line fit to the remaining genes, showing a 2.9-fold enrichment in fpkm values for all untreated transcripts. An R^2^ coefficient of 0.99 for this regression line indicates strong consistency between replicates with minimal off-target effects (Figure 2C).

To confirm that our depletion was specific to only the targeted mitochondrial sequences, we calculated the changes in fpkm values across all genes in the treated and untreated samples and identified those genes that were significantly diminished (> 2 standard deviations) relative to their control values. To overcome issues with stochastic variation at low gene counts/fpkm, we eliminated those genes that, between the three technical replicates at each Cas9 concentration, showed standard deviations in fpkm values greater than 50% of the mean. All of the genes meeting this criterion were present at less than 15 fpkm. Of the remaining genes, only one non-targeted human gene, MT-RNR2-L12, showed significant depletion when compared to the un-treated samples (Figure 2C). MT-RNR2-L12 is a pseudogene and shares over 90% sequence identity with a portion of the 16S mitochondrial rRNA gene. Out of the 23 sgRNA sites within the homologous region, 16 of them retain intact PAM sites in MT-RNR2-L12. Of these, seven have perfectly matching 20mer sgRNA target sites, and the remaining nine each have between one and four mutations (see Supplemental Figure 2). Depletion of this gene is therefore an expected consequence of our sgRNA choices.

### Reduction of Unwanted Abundant Sequences in CSF Samples

We next tested the utility of our method when applied to clinically relevant samples. In the case of pathogen detection in patient samples, the microbial transcripts are typically low in number and become greatly outnumbered by human host sequences. As a result, sequencing depth must be drastically increased to confidently detect such small minority sequence populations. We reasoned that depletion of unwanted high-abundance sequences from patient libraries could result in increased representation of pathogen-specific sequence reads. We thus integrated the DASH method with our in-house metagenomic deep sequencing diagnostic pipeline for patients with meningeal inflammation (i.e., meningitis) or brain inflammation (i.e., encephalitis) likely due to an infectious agent or pathogen. Figure 3 and Table 1 summarize the results of this analysis. In the case of a patient with meningoencephalitis whose CSF was previously shown to be infected with the amoeba *Balamuthia mandrillaris* [22], diagnosis was originally made by identification of a small fraction (<0.1%) of reads aligning to specific regions of the *B. mandrillaris* 16S mitochondrial gene. After DASH treatment, human mitochondrial 12S and 16S genes were reduced by an order of magnitude, and sequencing coverage of the *B. mandrillaris* 16S fragment increased 2.9-fold. Notably, *B. mandrillaris* is a eukaryotic organism, yet depletion of the human 16S gene by DASH did not have off target effects on the 16S *B. mandrillaris* mitochondrial gene. Similarly, patient CSF samples with confirmed *Taenia solium* (pork tapeworm) and *Cryptococcus neoformans* (fungus) infections showed 3.9 and 2-fold increases in coverage of the 18S genes of *T. solium* and *C. neoformans,* respectively, the detection of which was crucial in the initial diagnoses. The observed increases in relative signal can be translated into either a sequencing cost savings or a higher sensitivity that may be useful clinically for earlier detection of infections.

**Figure 3.**
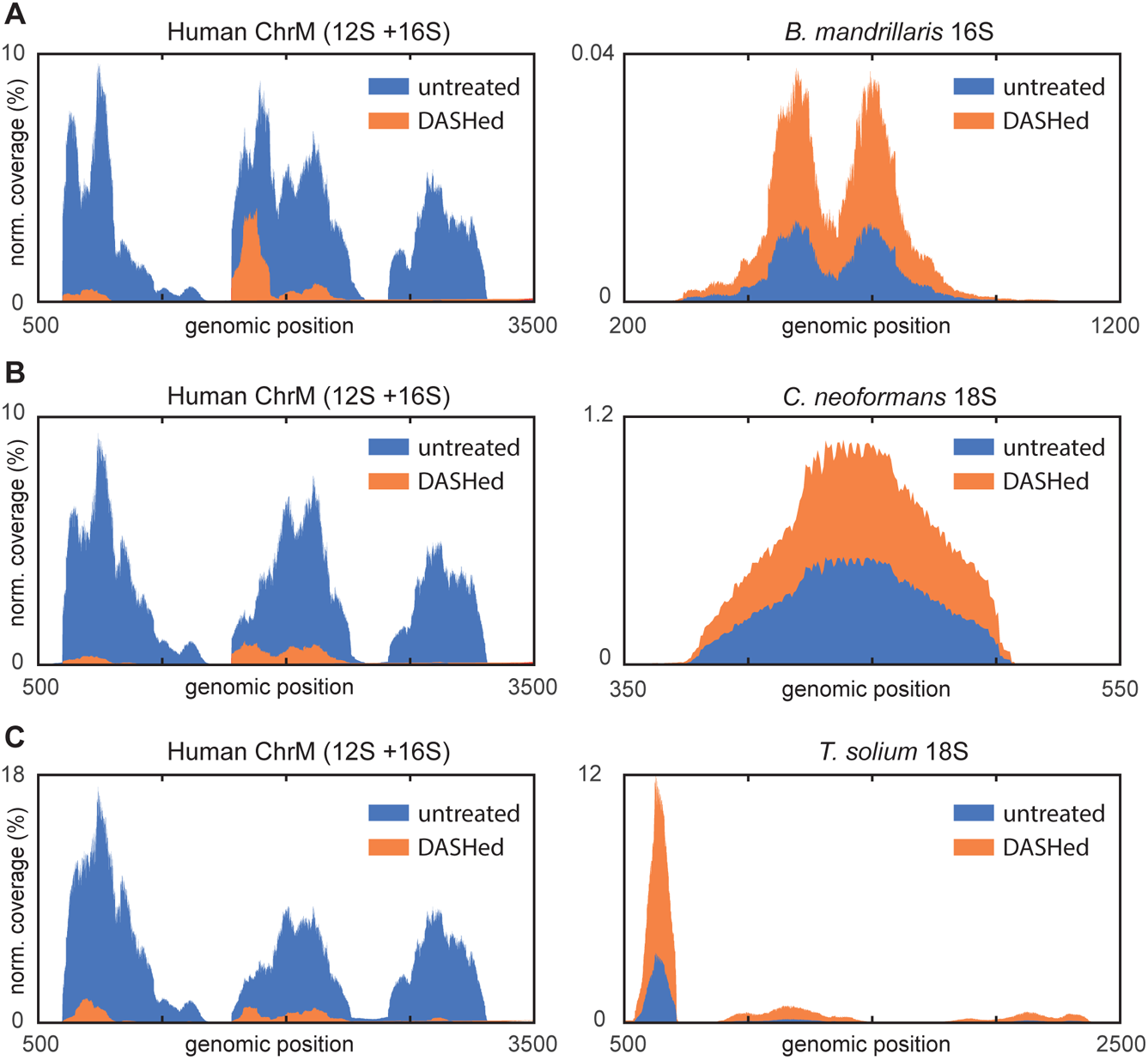
Normalized coverage plots of DASH-treated (orange) and untreated (blue) libraries generated from patient cerebrospinal fluid (CSF) samples with confirmed infections. Targeted mitochondrial rRNA genes (left) and representative genes for pathogen diagnosis (right) are depicted for the following: A) *Balamuthia mandrillaris,* B) *Cryptococcus neoformans,* C) *Taenia solium.* Across all cases, the DASH technique significantly reduced the coverage of human 12S and 16S genes by an average of 9.2-fold while increasing the coverage depth for pathogenic sequences by an average 2.93-fold. See Table 1 for relevant data.

### Reduction of Wild-type Background for Detection of the *KRAS* G12D (c.35G>A) Mutation in Human Cancer Samples

Specific driver mutations known to promote cancer evolution and at times to make up the genetic definition of malignant subtypes are important for diagnosis and targeted therapeutics (i.e. precision medicine). In complex samples isolated from biopsies or cell-free body fluids such as plasma, wild-type DNA sequences often overwhelm the signal from mutant DNA, making the application of traditional Sanger sequencing challenging [2, 3, 23]. For NGS, detection of minority alleles requires additional sequencing depth and therefore increases cost. We reasoned that the DASH technique could be applied to increase mutation detection from a PCR amplicon derived from a patient sample. We chose to focus on depletion of the wild-type allele of *KRAS* at the glycine 12 position, a hotspot of frequent driver mutations across a variety of malignancies [24–26]. This is an ideal site for DASH, because all codons encoding the wild-type glycine residue contain a PAM site (NGG), while any mutation that alters that residue (e.g., c.35G>A, p.G12D) ablates the PAM site and is thus uncleavable by Cas9 (see Figure 4A). This will be true of any mutation that changes a glycine (codons GGA, GGC, GGG, and GGT) or a proline (codons CCA, CCC, CCG, and CCT) to any other amino acid. Furthermore, it is relevant to the ubiquitous C>T nucleotide change found in germline mutations as well as somatic cancer mutations [27]. Targeting of other mutations will likely be possible in the near future with reengineered CRISPR nucleases or those that come from alternative species and have different PAM site specificities [28, 29].

**Figure 4.**
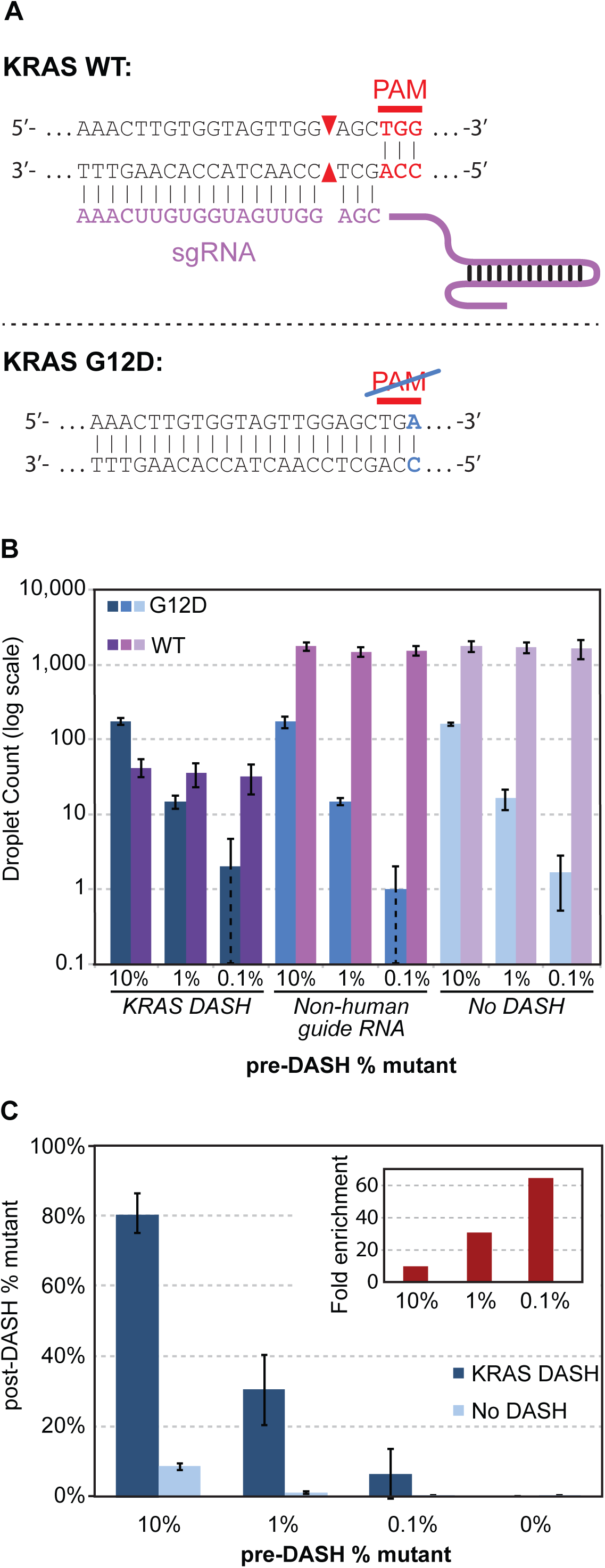
A) DASH is used to selectively deplete one allele while keeping the other intact. An sgRNA in conjunction with Cas9 targets a wild-type *KRAS* sequence. However, since the G12D (c.35G>A) mutation disrupts the PAM site, Cas9 does not efficiently cleave the mutant *KRAS* sequence. Subsequent amplification of all alleles using flanking primers, as in the case of digital PCR, Sanger sequencing, or high-throughput sequencing is only effective for non-cleaved and mutant sites. B) Three human genomic DNA samples with varying ratios of wild-type to mutant (G12D) *KRAS* were treated either with KRAS-targeted DASH, a non-human control DASH, or no DASH. Counts of intact wild-type and G12D sequences were then measured by droplet digital PCR (ddPCR). C) Same data as in B, presented as percentage of mutant sequences detected. Inset shows fold enrichment of the percentage of mutant sequences with KRAS-targeted DASH versus no DASH. For both B and C, values and error bars are the average and standard deviation, respectively, of three independent experiments.

The sequence of the sgRNA designed to target the *KRAS* G12D PAM site is listed in Supplemental Table 1, as is the non-human sequence used for the negative control sgRNA. Both were transcribed from a DNA template by T7 RNA polymerase, purified, and complexed with Cas9 as described in the Methods section. Samples were prepared by mixing sheared genomic DNA from a healthy individual (with wild-type *KRAS* genotype confirmed with digital PCR) and *KRAS* G12D genomic DNA to achieve mutant to wild type allelic ratios of 1:10, 1:100, and 1:1,000, and 0:1. For each mixture, 25 ng of a DNA mixture was incubated with 25 nM Cas9 pre-complexed with 25 nM of sgRNA targeting *KRAS* G12D. This concentration is high relative to the concentration of target molecules, but empirically we found it to be the most efficient ratio. We hypothesize that this may be due to non-cleaving Cas9 interactions with the rest of the human genome [21], which effectively reduce the Cas9 concentration at the cleavage site.

Samples were subsequently heated to 95°C for 15 minutes in a thermocycler to deactivate Cas9 (Methods). Droplet digital PCR (ddPCR) was used to count wild-type and mutant alleles using the primers and TaqMan probes depicted in Figure 4A and described in the Methods section. All samples were processed in triplicate. Samples incubated with or without Cas9 complexed to a non-human sgRNA target show the expected percentages of mutant allele: approximately 10%, 1%, and 0.1% for the 1:10, 1:100, and 1:1,000 initial mixtures respectively (Figure 4B). With addition of Cas9 targeted to *KRAS*, the wild-type allele count drops nearly two orders of magnitude (purple bars in Figure 4B), while virtually no change is observed in number of mutant alleles (blue bars). This confirms the high specificity of Cas9 for the NGG of the PAM site.

With the addition of DASH targeted to *KRAS* G12, the percentage of mutant allele jumps from 10% to 81%, from 1% to 30%, and from 0.1% to 6% (Figure 4C). This corresponds to 10-fold, 31-fold and 65-fold representational increases for the mutant allele, respectively. As expected, there was virtually no detection of mutant alleles in the wild type-only samples both with and without DASH treatment (one droplet in one of three no DASH wild type-only samples).

## Discussion

In this paper we have introduced DASH, a technique that leverages *in vitro* Cas9 ribonucleoprotein (RNP) activity to deplete specific unwanted high-abundance sequences which results in the enrichment of rare and less abundant sequences in NGS libraries or amplicon pools.

While the procedure may be easily generalized, we developed DASH to address current limitations in metagenomic pathogen detection and discovery, where the sequence abundance of an etiologic agent may be present as a minuscule fraction of the total. For example, infectious encephalitis is a syndrome caused by well over 100 pathogens ranging from viruses, fungi, bacteria and parasites. Because of the sheer number of diagnostic possibilities and the typically low pathogen load present in cerebrospinal fluid (CSF), more than half of encephalitis patients never have an etiologic agent identified [30]. We have demonstrated that NGS is a powerful tool for identifying infections, but as the *B. mandrillaris* meningoencephalitis case demonstrates, the vast majority of sequence reads are “wasted” resequencing high abundance human transcripts. In this case, we have shown that DASH depletes with incredible specificity the small number of human RNA transcripts that comprise the bulk of the NGS library, thereby lowering the required sequencing depth to detect non-human sequences and enriching the proportion of non-human (*Balamuthia*) reads in the metagenomic dataset.

In the case of infectious agents, it is possible to directly enrich rare sequences by hybridization to DNA microarrays [31] or beads [12]. However, these approaches rely on sequence similarity between the target and the probe and therefore may miss highly divergent or unanticipated species. Furthermore, the complexity and cost of these approaches will continue to increase with the known spectrum of possible agents or targets. In contrast, the identity and abundance of unwanted sequences in most human tissues and sample types has been well described in scores of previous transcriptome profiling projects [20], and therefore optimized collections of sgRNAs for DASH depletion are likely to remain stable.

DASH may also enhance the detection of rare mutant alleles that are important for liquid biopsy cancer diagnostics. Allelic depletion with DASH increases the signal (oncogenic mutant allele) to noise (wildtype allele) by more than 60 fold when studying the *KRAS* hotspot mutant p.G12D. With the rapidly growing number of oncologic therapies that target particular cancer mutations, sensitive and noninvasive techniques for cancer allele detection are increasingly relevant for optimizing the care of these patients [23]. These same techniques are also becoming increasingly important for diagnosis of earlier stage (and generally more curable) cancers as well as the detection of cancer recurrence without needing to re-biopsy the patient [2, 14, 32, 33, 33, 34].

The potential applications of DASH are manifold. Currently, DASH can be customized to deplete any set of defined PAM-adjacent sequences by designing specific libraries of sgRNAs. Given the popularity and promise of CRISPR technologies, we anticipate the adaptation and/or engineering of CRISPR-associated nucleases with more diverse PAM sites [28, 29, 35]. A portfolio of next-generation Cas9-like nucleases would further enable DASH to deplete large and diverse numbers of arbitrarily selected alleles across the genome without constraint. We envision that DASH will be immediately useful for the development of non-invasive diagnostic tools, with applications to low input samples or cell-free DNA, RNA, or methylation targets in body fluids [4, 6, 32, 34, 36, 37].

Many other NGS applications could also benefit from depletion of specific sequences, including hemoglobin mRNA depletion for RNA-Seq of blood samples [38] and tRNA depletion for ribosome profiling studies. Depletion of pseudogenes or otherwise homologous sequences by small but consistent differences in sequences is also theoretically possible, and may serve to remove ambiguities in clinical high-throughput sequencing. Using DASH to enrich for minority variations in microbial samples may enable early discovery of pathogen drug resistance. Similarly, the application of DASH to the analysis of cell-free DNA may augment our ability to detect early markers of drug resistance in tumors [23].

## Conclusions

Here, we have demonstrated the broad utility of DASH to enhance molecular signals in diagnostics and its potential to serve as an adaptable tool in basic science research. While the degree of regional depletion of mitochondrial rRNA was sufficient for our application, the depletion parameters were not maximized: we used only 54 sgRNA target sites out of about 250 possible S. *pyogenes* Cas9 sgRNA candidates in the targeted mitochondrial region. Future studies will explore the upper limit of this system while elucidating the most effective sgRNA and CRISPR-associated nuclease selections, which will likely differ based on target and application. Irrespective, depletion of unwanted sequences by DASH is highly generalizable and may effectively lower costs and increase meaningful output across a broad range of sequence-based approaches.

## Methods

### Generation of cDNA from HeLa cell line and clinical samples

CSF samples were collected under the approval of the institutional review boards of the University of California San Francisco and San Francisco General Hospital. Samples were processed for high-throughput sequencing as previously described [1, 22]. Briefly, amplified cDNAs were made from randomly primed total RNA extracted from 250uL of CSF or 250 pg of HeLa RNA using the NuGEN Ovation v.2 kit (NuGEN, San Carlos, CA) for low nucleic acid content samples. A Nextera protocol (Illumina, San Diego, CA) was used to add on a partial sequencing adapter on both sides.

### In vitro preparation of the CRISPR/Cas9 complex

The Cas9 expression vector, containing an N-terminal MBP tag and C-terminal mCherry, was kindly provided by Dr. Jennifer Doudna. The protein was expressed in BL21 Rosetta cells for three hours at 18°C. Cells were pelleted and frozen. Upon thawing, cells from a 4 L culture prep were resuspended in 50 mL of lysis buffer (50 mM sodium phosphate pH 6.5, 350 mM NaCl, 1 mM TCEP, 10% glycerol) supplemented with 0.5 mM EDTA, 1 μM PMSF, and a single Roche complete EDTA-free protease inhibitor tablet (Roche Diagnostics, Indianapolis, IN) and passed through an HC-8000 homogenizer (Microfluidics, Westwood, MA) five times. The lysate was clarified by centrifugation at 20,000 rpm for 45 minutes at 4°C and then filtered through a 0.22 μm vacuum filtration unit. The filtered lysate was loaded onto three 5 mL HiTrap Heparin HP columns (GE Healthcare, Little Chalfont, UK) arranged in series on a GE AKTA Pure system. The columns were washed extensively with lysis buffer, and the protein was eluted with a gradient of lysis buffer to buffer B (lysis buffer supplemented with NaCl up to 1.5M). The resulting fractions were analyzed by Coomassie gel, and those containing Cas9 (centered around the point on the gradient corresponding to 750 mM NaCl) were combined and concentrated down to a volume of 1 mL using 50K MWCO Amicon Ultra-15 Centrifugal Filter Units (EMD Millipore, Billerica, MA) and then fed through a 0.22 µm syringe filter. Using the AKTA Pure, the 1 mL of filtered protein solution was then injected onto a HiLoad 16/600 Superdex 200 size exclusion column (GE Healthcare, Little Chalfont, UK) pre-equilibrated with buffer C (lysis buffer supplemented with NaCl up to 750 mM). Resulting fractions were again analyzed by Coomassie gel, and those containing purified Cas9 were combined, concentrated, supplemented with glycerol up to a final concentration of 50%, and frozen at − 80°C until use. Protein concentration was determined by BCA assay. Yield was approximately 80 mg from 4 L of bacterial culture.

sgRNA target sites were selected as described in the main text. DNA templates for sgRNAs based on an optimized scaffold [39] were made with a similar method to that described in [40]For each chosen target, a 60mer oligo was purchased including the 18-base T7 transcription start site, the targeted 20mer, and the first 22 bases of the tracr RNA (5’-TAATACGACTCACTATAG NNNNNNNNNNNNNNNNNN NNGTTTAAGAGCTATGCTGGAAAC-3’). This was mixed with a 90mer representing the 3’ end of the sgRNA on the opposite strand (5’-AAAAAAAGCACCGACTCGGTGCCACTTTTTCAAGTTGATAACGGACTAGCCTTATTTAAACTTGCTATGCTGTTTCCAGCATAGCTCTTA-3’). DNA templates for T7 sgRNA transcription were then assembled and amplified with a single PCR reaction using primers 5’-TAATACGACTCACTATAG-3’ and 5’-AAAAAAAGCACCGACTCGGTGC-3’. The resulting 131 base pair (bp) transcription templates, with the sequence 5’-TAATACGACTCACTATAG NNNNNNNNNNNNNNNNNNNNGTTTAAGAG CTATGCTGGAAACAGCATAGCAAGTTTAAATAAGGCTAGTCCGTTATCAACTTGAAAAAGTGGCACCGAGTCGGTGCTTTTTTT-3’, were pooled (for the mitochondrial rRNA library), or transcribed separately (for the *KRAS* experiments). All oligos were purchased from IDT (Integrated DNA Technologies, Coralville, IA).

Transcription was performed using custom-made T7 RNA polymerase (RNAP) [41, 42] In each 50 μL reaction, 300 ng of DNA template was mixed with T7 RNAP (final concentration 8 ng/μL), buffer (final concentrations of 40 mM Tris pH 8.0, 20 mM MgCl_2_, 5 mM DTT, and 2 mM spermidine), and Ambion brand NTPs (ThermoFisher Scientific, Waltham, MA) (final concentration 1 mM each ATP, CTP, GTP and UTP), and incubated at 37°C for 4 hours. Typical yields were 2-20 μg of RNA. sgRNAs were purified with a Zymo RNA Clean & Concentrator-5 kit (Zymo Research, Irvine, CA), aliquoted, stored at −80°C, and used only a single time after thawing.

### CRISPR/Cas9 treatment

To form the ribonucleoprotein (RNP) complex, Cas9 and the sgRNAs were mixed at the desired ratio with Cas9 buffer (final concentrations of 50 mM Tris pH 8.0, 100 mM NaCl, 10 mM MgCl_2_, and 1 mM TCEP), and incubated at 37°C for 10 minutes. This complex was then mixed with the desired amount of sample cDNA in a total of 20 μL, again in the presence of Cas9 buffer, and incubated for 2 hours at 37°C.

Since Cas9 has high nonspecific affinity for DNA [21] it was necessary to disable and remove the Cas9 before continuing. For the rRNA depletion samples, 1 μL (at >600 mAU/mL) of Proteinase K (Qiagen, Hilden, Germany) was added to each sample which was then incubated for an additional 15 minutes at 37°C. Samples were then expanded to a volume of 100 μL and purified with three phenol:chloroform:isoamyl alcohol extractions followed by one chloroform extraction in 2mL Phase-lock Heavy tubes (5prime, Hilden, Germany). 10 μL of 3M sodium acetate pH 5.5, 3 μL of linear acrylamide and 226 μL of 100% ethanol were added to the 100 μL aqueous phase of each sample. Samples were cooled on ice for 30 minutes. DNA was then pelleted at 4°C for 45 minutes, washed once with 70% ethanol, dried at room temperature and resuspended in 10 μL water.

In the case of the *KRAS* samples, Cas9 was disabled by heating the sample at 95°C for 15 minutes in a thermocycler and then removed by purifying the sample with a Zymo DNA Clean & Concentrator-5 kit (Zymo Research, Irvine, CA).

### High-throughput sequencing and Analysis of sequencing data

Tagmented samples with and without DASH treatment underwent 10-12 cycles of additional amplification (Kapa Amplification Kit, Kapa Biosystems, Wilmington, MA, USA) with dual-indexing primers. A BluePippin instrument (Sage Science, Beverly, MA, USA) was used to extract DNA between 360-540 bp. Sequencing libraries were purified using the Zymo DNA Clean & Concentrator-5 kit and amplified again on an Opticon qPCR machine (MJ Research, Waltham, MA, USA) using a Kapa Library Amplification Kit until the exponential portion of the qPCR signal was found. Sequencing libraries were then pooled and re-quantified with a droplet digital PCR (ddPCR) Library Quantification Kit (Bio-Rad, Hercules, CA). Sequencing was performed on portions of one lane in an Illumina HiSeq 4000 instrument using 135 bp-end sequencing.

All reads were quality filtered using PriceSeqFilter v1.2 [43] such that only read pairs with less than 5 ambiguous base calls (defined as N’s or positions with < 95% confidence based on Phred score) were retained. Filtered reads were aligned to the hg38 build of the human genome using the STAR aligner (v 2.4.2a) [44]. Gene counts were generated using the Gencode v23 primary annotations. Pathogen-specific alignments to 16S and 18S sequences were accomplished using Bowtie2 [45]). Per-nucleotide coverage was calculated from alignment (SAM/BAM) files using the SAMtools suite [46] and analyzed with custom iPython [47] scripts utilizing the Pandas data package. Plots were generated with Matplotlib [48].

### Digital PCR of *KRAS* mutant DNA

*KRAS* wild-type DNA was obtained from a healthy consenting volunteer. The sample sat until cell separation occurred, and DNA was extracted from the buffy coat with the QIAamp Blood Mini Kit (Qiagen, Hilden, Germany). *KRAS* G12D genomic DNA from the human leukemia cell line CCRF-CEM was purchased from ATCC (Manassas, VA). All DNA was sheared to an average of 800 bp using a Covaris M220 (Covaris, Woburn, USA) following the manufacturer’s recommended settings. Cas9 reactions occurred as described above.

A primer/probe pair was designed with Primer3 [49, 50] targeting the relatively common *KRAS* G12D (c.35G>A) mutation. Reactions were themocycled according to manufacturer protocols using a 2-step PCR. An ideal 62°C annealing/extension temperature was determined by a gradient experiment to ensure proper separation of FAM and HEX signals. The PCR primers and probes used were as follows (purchased from IDT): Forward: 5’-TAGCTGTATCGTCAAGGCAC – 3’, Reverse: 5’ – GGCCTGCTGAAAATGACTGA – 3’, wild-type probe: 5’ – /5HEX/TGCCTACGC/ZEN/CA<C>CAGCTCCA/3IABkFQ/ – 3’, mutant probe: 5’ – /56-FAM/TGCCTACGC/ZEN/CA<T>CAGCTCCA/3IABkFQ/ – 3’, with <> denoting the mutant base location, 5HEX and 56-FAM denoting the HEX and FAM reporters, and ZEN and 3IABkFQ denoting the internal and 3’ quenchers. Original samples and those subjected to DASH were measured with the ddPCR assay on a Bio-Rad QX100 Droplet Digital PCR system (Bio-Rad, Hercules, CA), following the manufacturer’s instructions for droplet generation, PCR amplification, and droplet reading, and using best practices. Pure CCRF-CEM samples was approximately 30% G12D and 70% wild type; all calculations of starting mixtures were made based on this starting ratio.

## Acknowledgements

We thank Derek Bogdanoff and the Center for Advanced Technologies at UCSF for assistance with sequencing and help with equipment; Hannah Sample for assistance with sample collection; Dr. Joe Kliegman for HeLa RNA; Jennifer Mann for administrative support; Dr. Chong Park and the Innovative Genomics Initiative for help with sgRNA production protocols; Lara Pesce Ares for help with Cas9 purification; Drs. Sy Redding, Wilson Koh, and Charles Chiu for helpful discussions; and Drs. Jeffrey Gelfand, Luke Strnad and Niraj Shanbhag for patient referrals. We would also like to thank the patients and their families who volunteered to participate in our research program. This work was supported by the Sandler Foundation, the William K. Bowes, Jr. Foundation, Howard Hughes Medical Institute (to ED Crawford and JLD), and the National Center for Advancing Translational Sciences of the NIH [KL2TR000143] (to MRW). The contents of this paper are solely the responsibility of the authors and do not necessarily represent the official views of the NIH.

**List of abbreviations**
DASH: Depletion of Abundant Sequences by Hybridization
NGS: Next Generation Sequencing
CRISPR: Clustered Regularly Interspaced Short Palindromic Repeats
Cas: CRISPR-associated
sgRNA: single guide RNA
CSF: Cerebrospinal Fluid
ddPCR: droplet digital PCR

## Competing Interests

WG is a consultant for Pacgeno and holds patents relevant to microfluidics and non-invasive prenatal diagnostics. ED Crawford, BDO, MRW, ED Chow, HR and JLD have no competing interests.

## Authors’ contributions

WG and EDC1 (Crawford) contributed equally to this work and are co-first authors. WG, EDC1, BDO, MRW, HR, EDC2 (Chow), and JLD conceived of the project. EDC1, WG, MRW, and JLD designed and performed the experiments. MRW oversaw the encephalitis study and prepared the original NGS libraries used to identify pathogens and determine high-abundance sequences. EDC1 produced and optimized the *in vitro* Cas9 and sgRNA system and directed its usage. EDC2 performed sequencing. BDO, WG, EDC1, JLD analyzed the data. EDC1, WG, BDO and JLD drafted the manuscript with input and edits from all other authors.

## Supplemental Figures/Table/Attachments

Supplemental Table 1: List of all sgRNA sequences used in this paper.

Supplemental Figure 1: Depletion efficiency by Cas9 dosage.

Supplemental Figure 2: PAM sites in pseudogene.

Supplemental Figure 3: Scatter plots for patient samples.

**Supplemental Table 1:**
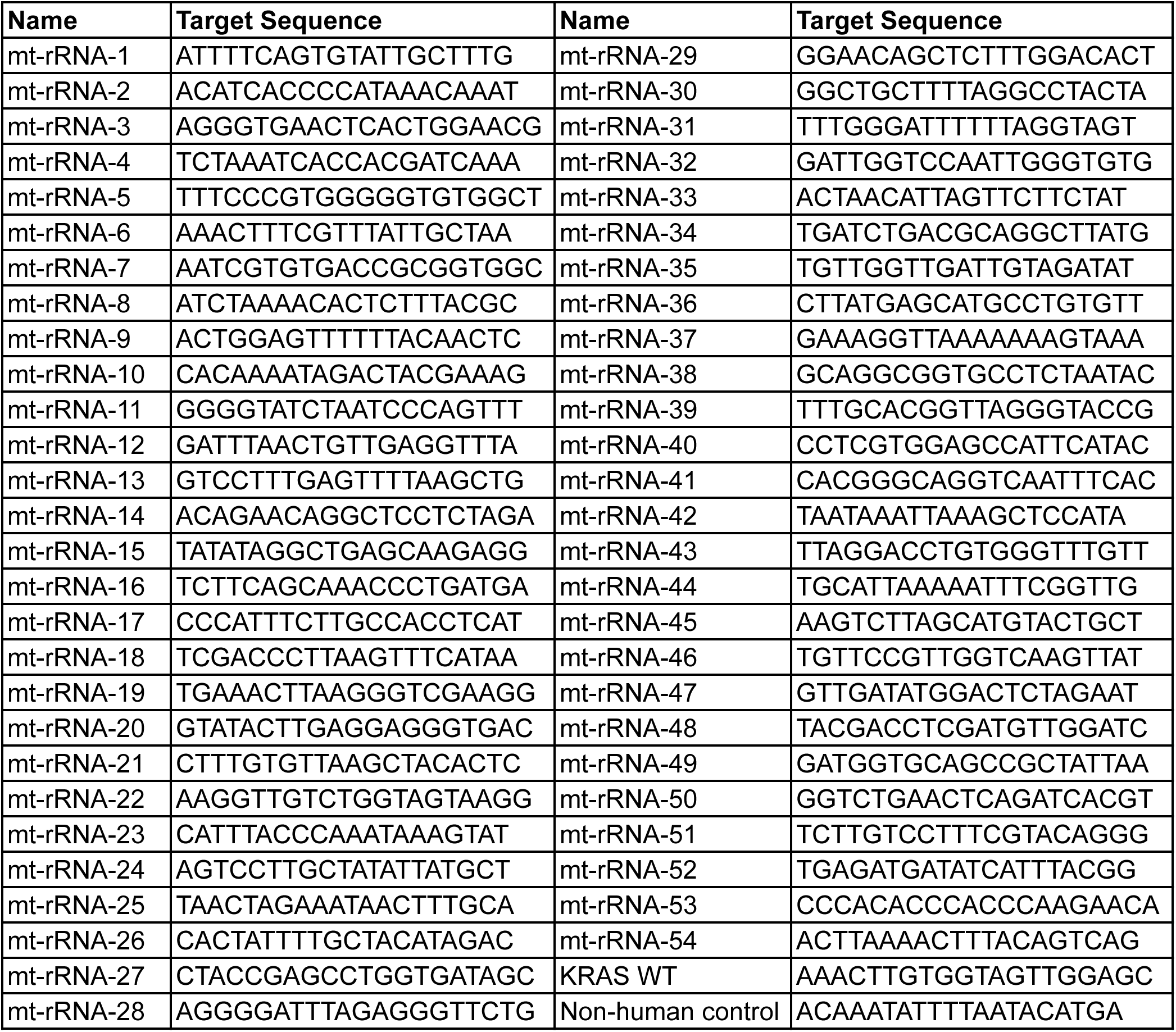
sgRNA Target sequences

**Supplemental Figure 1.**
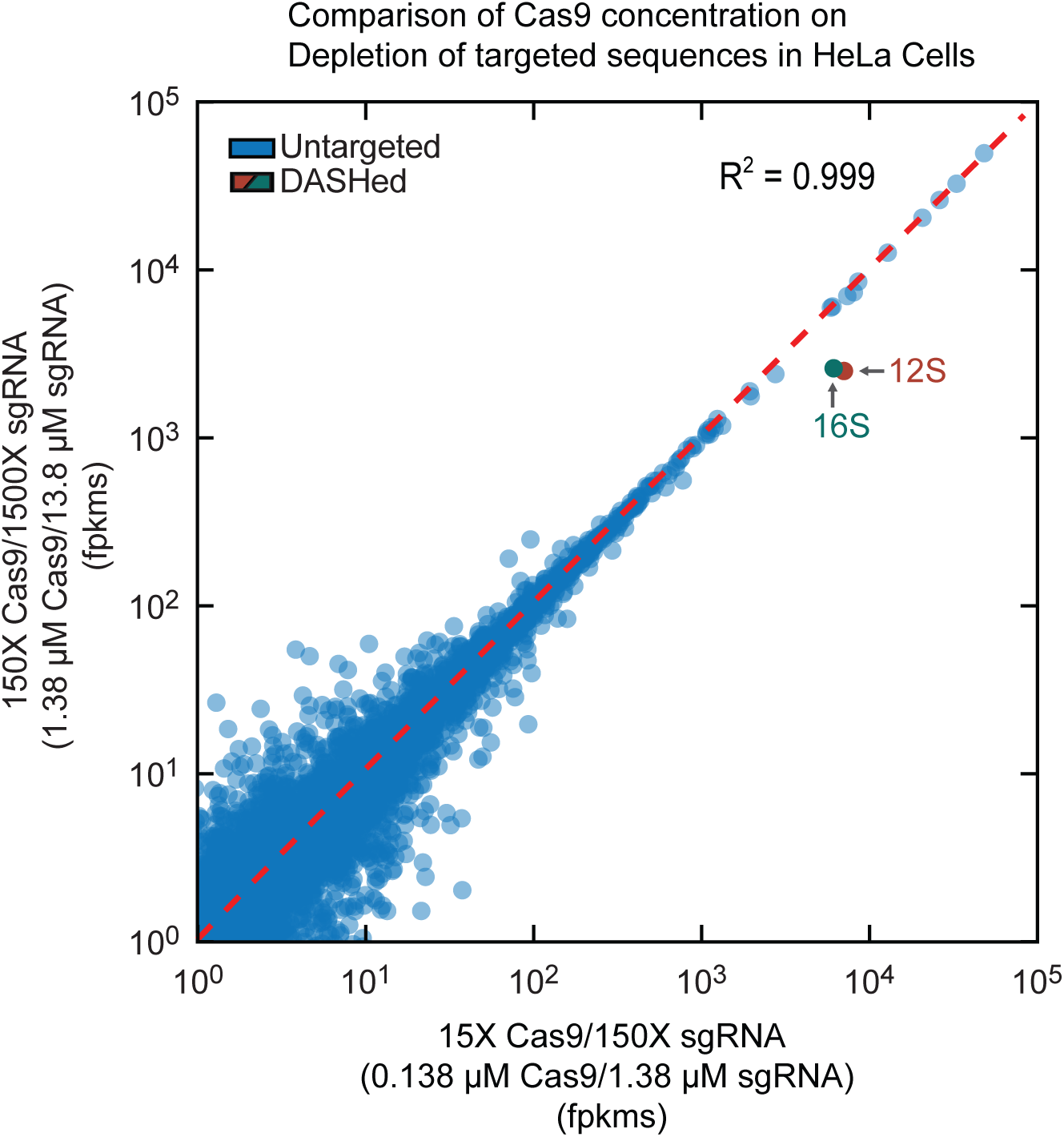
Scatterplot of log of fragments per kilobase of transcript per million mapped reads (log-fpkm) values per human gene comparing HeLa cells DASHed with two different Cas9/sgRNA concentrations.

**Supplemental Figure 2.**
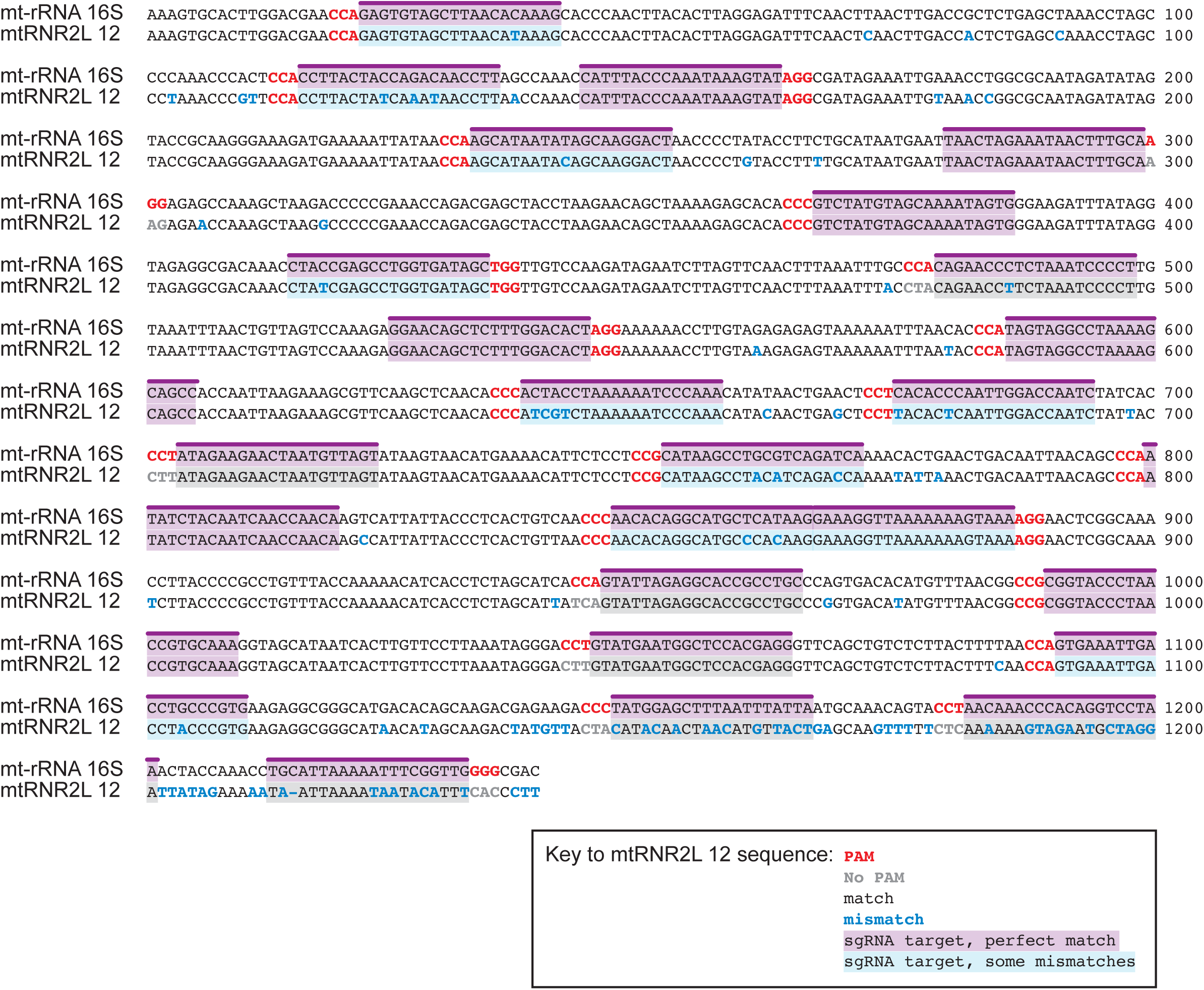
mt rRNA sgRNA target sites in the pseudogene mtRNR2L 12.

**Supplemental Figure 3.**
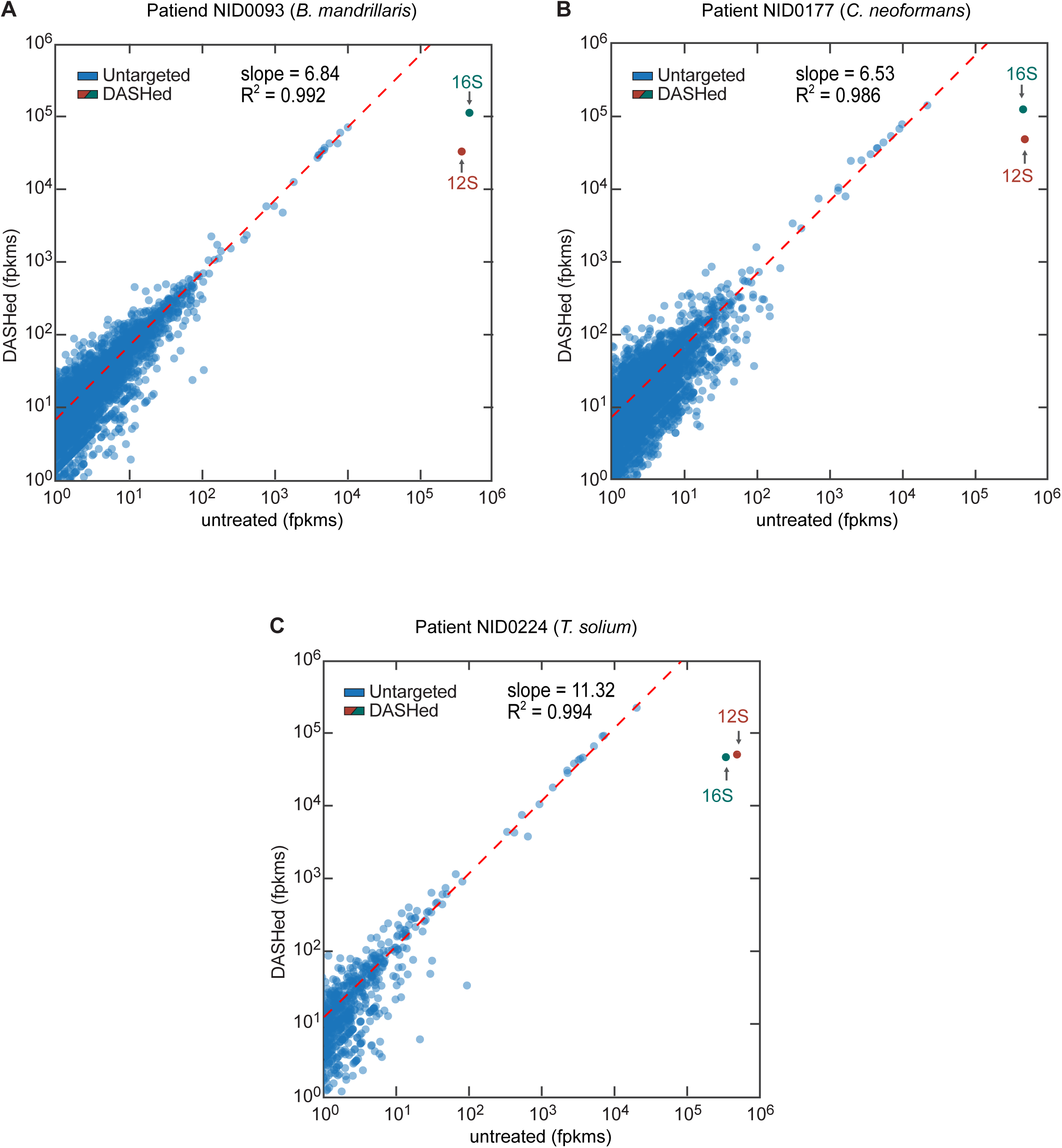
Scatterplots of the log of fragments per kilobase of transcript per million mapped reads (log-fpkm) values per human gene in the DASHed vs. untreated patient samples. The slopes of the regression lines (red) indicate the fold enrichment in reads mapped to untargeted transcripts. R-squared (R^2^) values of the regression lines indicate minimal off-target depletion.

